# SG33, a vaccine strain of myxoma virus with oncolytic potential, exploits macropinocytosis and clathrin-mediated endocytosis for entry into pancreatic cancer cells

**DOI:** 10.64898/2025.12.16.694613

**Authors:** Nikolaos Kontopoulos, Noémie Delenclos, Laetitia Ligat, Sokunthea Top, Franck Gallardo, Stephane Bertagnoli, Nelson Dusetti, Louis Buscail, Pierre Cordelier

## Abstract

Oncolytic viruses are being investigated as therapeutic agents in cancer, yet their mechanisms of entry into tumor cells remain incompletely understood. We previously showed that SG33, a veterinary vaccinal strain derived from a pathogenous myxoma virus, displays oncolytic activity in preclinical models of pancreatic ductal adenocarcinoma (PDAC). Here, we investigated the entry pathways of SG33 into primary PDAC-derived cultures. We found that macropinocytosis, an endocytic process frequently upregulated in PDAC, contributes to SG33 uptake. Moreover, SG33 infection itself induced macropinocytosis in a subset of primary PDAC cultures. Mechanistic studies revealed that phosphatidylserine exposed on the viral envelope promotes SG33 internalization through apoptotic mimicry. In PDAC cultures lacking detectable macropinocytosis, SG33 employed clathrin-mediated endocytosis as an alternative entry route. These findings provide the first insights into the entry mechanisms of SG33 into PDAC-derived cells and indicate that this virus can utilize distinct endocytic pathways depending on the cellular context.

**Importance:** Pancreatic ductal adenocarcinoma (PDAC) remains one of the most lethal cancers, with limited therapeutic options. Oncolytic virotherapy is emerging as a promising strategy to overcome treatment resistance, yet the mechanisms by which candidate viruses enter cancer cells remain poorly defined. Here, we characterize the entry route of SG33, a derivative of myxoma virus with potent oncolytic activity in PDAC models. We show that SG33 exploits multiple endocytic pathways, including macropinocytosis (MPC), induces MPC through apoptotic mimicry, and also uses clathrin-mediated endocytosis (CME). These findings provide the first evidence that SG33 has evolved a flexible cell entry strategy, which could enhance its efficacy in the heterogeneous context of PDAC. Understanding the molecular determinants of viral entry is essential for the rational design of improved oncolytic virotherapies. Our study is aligned with this objective and highlights SG33 as a promising candidate to expand the toolbox of virotherapeutic agents against aggressive cancers.

## INTRODUCTION

Oncolytic viruses (OVs) have gained significant attention in biomedicine due to their ability to selectively infect and lyse cancer cells while sparing normal cells(1). In addition, OVs can stimulate immune responses by exposing tumor-associated antigens, thereby converting immunologically “cold” tumors into “hot” tumors (1). This dual activity, combining direct oncolysis with immune activation, has made OVs attractive candidates for cancer therapy. Despite many OVs progressing to clinical trials, only two have been approved for clinical use (2, 3). The therapeutic activity of OVs can be limited by interpatient and intratumoral heterogeneity, as shown for the Minute Virus of Mice (MVM) which preferentially infects cancer cells with a mesenchymal phenotype (4). Physical barriers may also restrict viral penetration into tumors, and intrinsic cellular restriction factors against OVs are constitutively expressed in some cancer cells (5). Thus, characterization of viral tropism is critical to improve OV efficacy, particularly by identifying cancer cell subtypes that are more permissive to infection. Such studies may ultimately guide the development of more effective and tailored therapies for cancer.

A critical determinant of viral tropism, and ultimately of the efficacy of oncolytic virotherapy, is viral entry, which may occur either through direct fusion with the cell membrane or via endocytosis (6). Several endocytic pathways have been described: clathrin- and caveolin-mediated endocytosis typically internalize small volumes, whereas macropinocytosis (MPC) and phagocytosis are associated with the uptake of larger volumes (7). MPC is a highly regulated, growth factor-nduced and actin-dependent, endocytic pathway. Many viruses exploit MPC as a “Trojan horse” to enter host cells, evade antiviral responses, and ensure replication and propagation (8). MPC is frequently upregulated in cancers, including pancreatic ductal adenocarcinoma (PDAC) (9), a disease projected to become the second leading cause of cancer-related death worldwide (10). Current treatments, mainly chemotherapy, provide only limited benefit, extending survival by a few weeks to months (11, 12). Although PDAC has been proposed as a candidate for oncolytic virotherapy, no OV has yet gained approval or demonstrated success in phase III clinical trials (13). In PDAC, MPC has emerged as an important mechanism by which cancer cells acquire nutrients to sustain high metabolic demands, particularly under adverse conditions (14, 15). This process can occur in two modes, basal and inducible, allowing cancer cells to adapt to changing microenvironmental cues(16).

Among OVs, Myxoma virus (MYXV) has shown promising activity in experimental models of PDAC (17, 18). MYXV is a large, enveloped poxvirus with a double-stranded DNA genome that replicates in the cytoplasm (19). It naturally infects European rabbits (*Oryctolagus cuniculus*), causing myxomatosis. Studies on vaccinia virus, a close relative of MYXV, demonstrated that this virus can exploit macropinocytosis (MPC) to enter cells (20), and Villa *et al.* reported that vaccinia virus and MYXV employ distinct endocytic pathways in human cancer cells (21). In contrast, Duteyrat *et al.* suggested that MYXV entry into permissive cells occurs via an endocytic process resembling MPC (22). Thus, the literature provides conflicting views on the entry mechanisms of MYXV.

We previously demonstrated that SG33, an attenuated MYXV strain originally developed as a veterinary vaccine to protect rabbits from myxomatosis (23), displays enhanced oncolytic activity compared to wild type strain (24). In addition, SG33 shows activity in primary cultures and experimental PDAC tumors (24). These features identify SG33 as a potential candidate for oncolytic virotherapy in PDAC. However, the mode of entry of SG33 into cancer cells remains unknown.

Here we determine whether MPC contributes to SG33 tropism in PDAC-derived cultures. We first assessed basal and inducible MPC levels in several pancreatic cancer cell lines and primary cultures. We then examined the impact of MPC inhibition on SG33 entry and replication. Our results show that SG33 entry and replication are impaired when MPC is inhibited in permissive primary PDAC cells. Furthermore, we found that SG33 induces MPC through an apoptotic mimicry strategy, and that it can also utilize clathrin-mediated endocytosis in PDAC cultures lacking detectable MPC activity. These findings provide the first insights into the entry mechanisms of SG33 in PDAC-derived cells and indicate that SG33 can employ multiple endocytic pathways depending on the cellular context.

## RESULTS

### Characterization of basal and inducible MPC levels in PDAC-derived cell models

MPC levels in PDAC cell lines have been reported (16, 27), but data on patient-derived culture models remain limited. To address this, we analyzed basal MPC activity in primary PDAC cultures (PDAC087T, PDAC091T, PDAC084T, PDAC012T, PDAC015T, PDAC072T, PDAC051T) and compared them with PDAC cell lines with known MPC activity (MIA PaCa-2, AsPC-1, BxPC-3). MPC activity was measured using uptake of 70-kDa FITC-dextran, with the specific inhibitor EIPA as control (28). Serum-starved PDAC087T cells were treated with increasing concentrations of EIPA and assayed for 70-kDa dextran uptake and proliferation. Treatment with 75 μM EIPA reduced dextran uptake by 77% ± 10 (p<0.05) with minimal effect on proliferation (Fig. S1A-C). This concentration was used for subsequent experiments (Fig. 1A-B). We next quantified FITC-dextran uptake in the PDAC cultures. MIA PaCa-2 and PDAC087T cells showed high MPC activity (mean 5.8 × 10e4 and 16 × 10e4 relative units (r.u.), respectively), whereas AsPC-1, BxPC-3, PDAC091T, PDAC084T, PDAC012T, PDAC015T, PDAC072T, and PDAC051T displayed lower activity (4.2 × 10e3 to 2.1 × 10e4 r.u.) (Fig. 1C bottom panel, 1D). Similar results were obtained with 10-kDa dextran (Fig. 1C, top panel, 1D). Together, these data define conditions for monitoring MPC and reveal marked heterogeneity among PDAC models, allowing classification into MPC-high (MIA PaCa-2, PDAC087T) and MPC-low groups (all others).

**Figure 1.**
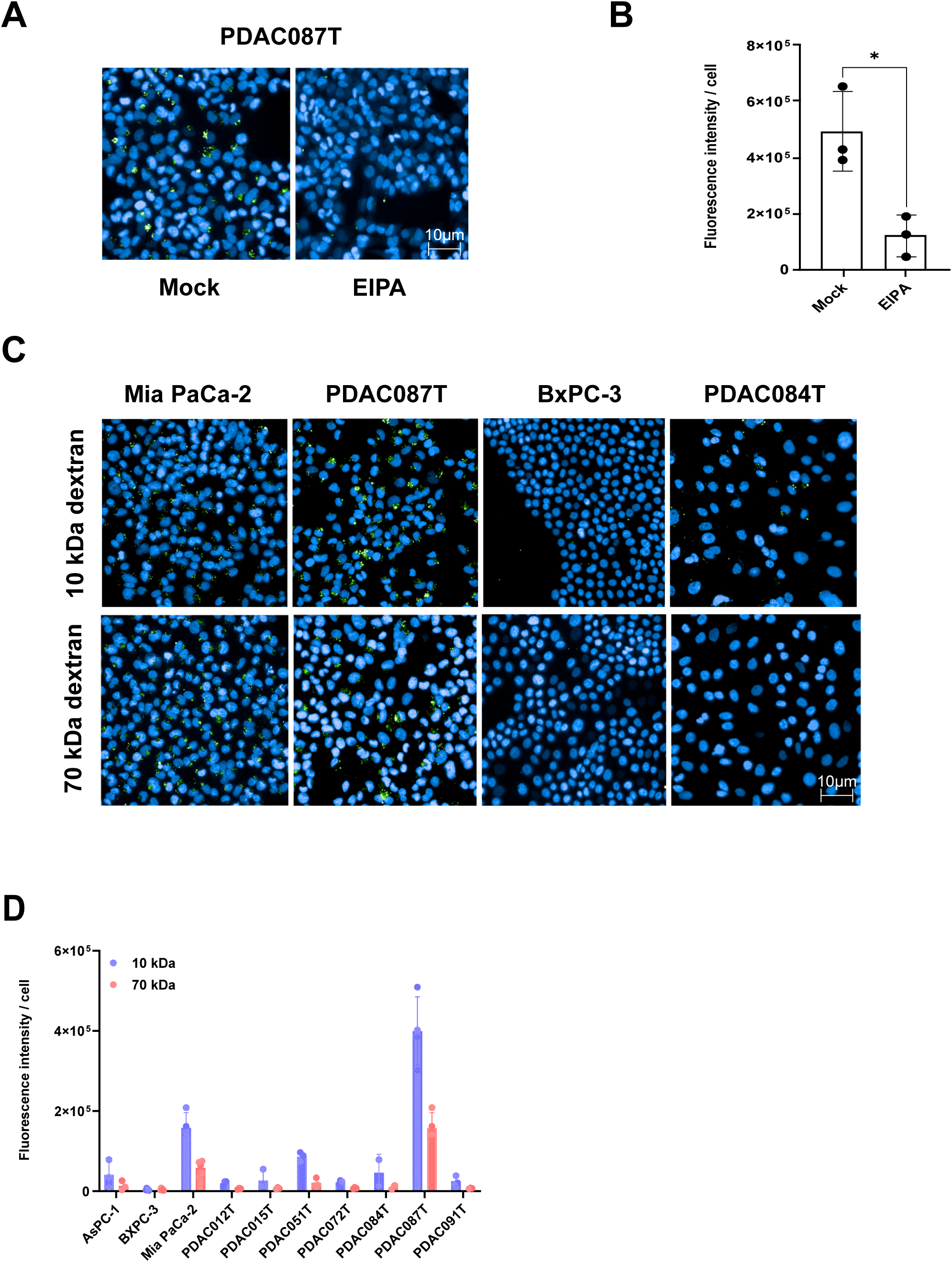
Basal macropinocytosis in PDAC cell lines and patient-derived cultures. Representative images (A) and quantification (B) of 70 kDa FITC-dextran (1 mg/mL) uptake in PDAC087T cells treated with EIPA (5-(N-ethyl-N-isopropyl) amiloride, 75 µM, 30 min). Control cells received DMSO. Results represent mean ± SD of three independent experiments performed in triplicate. Nuclei were stained with DAPI (blue). Representative images (C) and quantification (D) of 10 kDa (top) or 70 kDa (bottom) FITC-dextran (1 mg/mL) uptake in the indicated PDAC cells. 180 fields (∼25,000 cells) were analyzed

### Inducible macropinocytosis in PDAC primary cultures

In PDAC, MPC can be induced by growth factors or nutrient deprivation(16). To assess inducible MPC, primary pancreatic cultures were serum-starved and then treated with epidermal growth factor (EGF, 100 ng/ml) or cultured in glutamine-free medium, conditions previously reported to stimulate MPC in PDAC cells (25). In AsPC-1 cells, serum starvation followed by EGF treatment for 5 minutes increased 70-kDa FITC-dextran uptake 5.1-fold ± 0.2 (p<0.001, Fig. 2A, B). EIPA treatment inhibited EGF-mediated dextran uptake by 71% ± 9 (p<0.05, Fig. 2C, D), confirming MPC induction. Glutamine deprivation produced a similar effect, with a 4.8-fold ± 1.5 increase in dextran uptake (p<0.05, Fig. 2A, B). Western blot analysis of serum-starved AsPC-1 cells treated with EGF revealed phosphorylation of ERK1/2, EGFR and as well as increased PAK activation, a marker of inducible MPC (Supplementary Fig. S2A-D). Analysis of primary PDAC cultures showed that EGF increased dextran uptake in PDAC087T (+29% ± 2, p<0.005) and PDAC091T (+51% ± 1, p<0.05, Fig. 2E), with similar results under glutamine deprivation (Supp. Fig. S2E). These data demonstrate that PDAC primary cultures exhibit heterogeneous basal MPC levels and variable inducibility in response to growth factor stimulation or nutrient deprivation.

**Figure 2.**
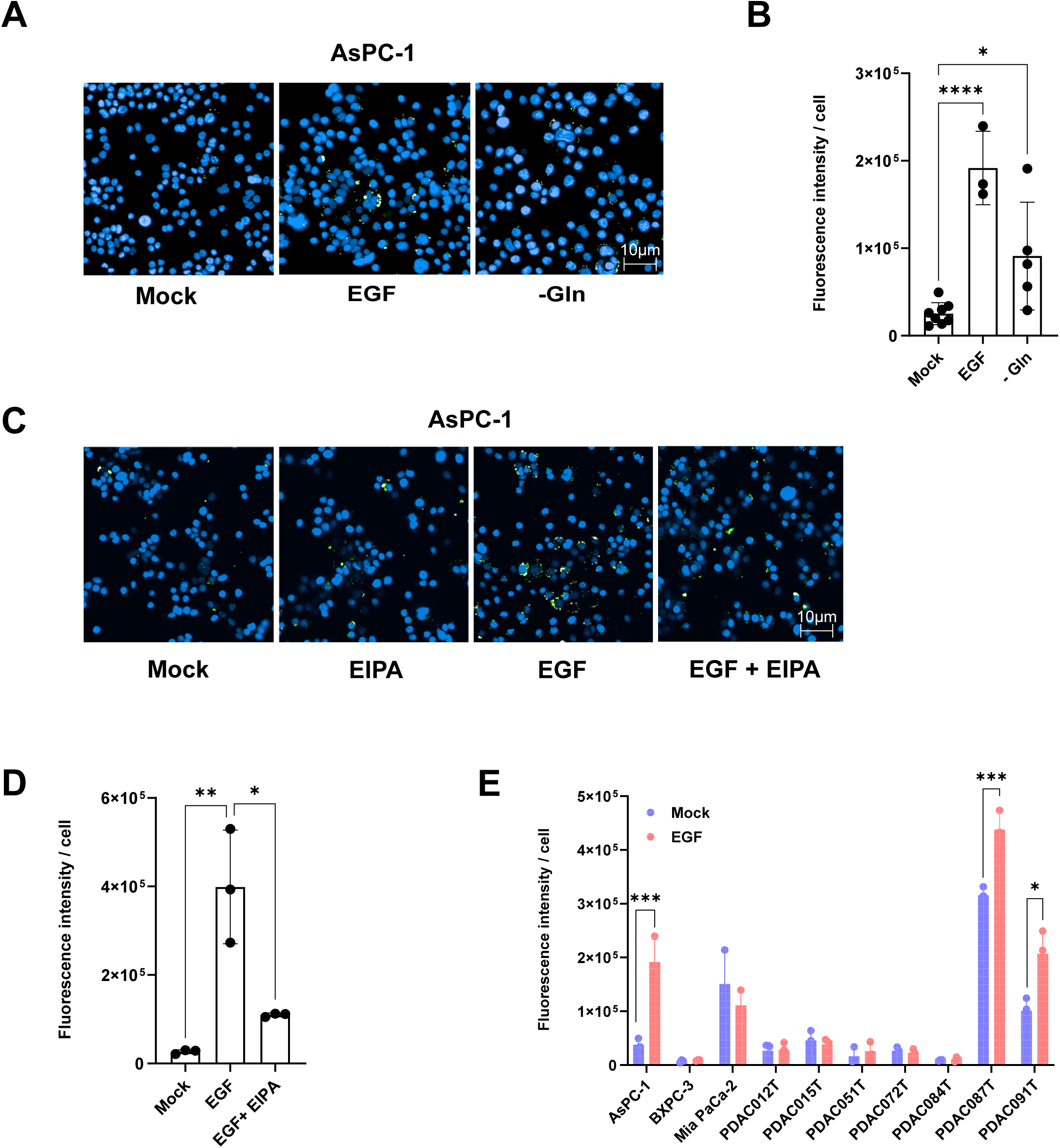
Inducible macropinocytosis in PDAC cell lines and patient-derived cultures. Representative images (A) and quantification (B) of 70 kDa FITC-dextran (1 mg/mL) uptake in AsPC-1 cells following treatment with EGF (100 ng/mL, 5 min) or glutamine starvation (-Gln, 18 h). Control cells were cultured in starvation medium supplemented with 2 mM glutamine. Results represent mean ± SD of at least three independent experiments performed in triplicate. Nuclei were stained with DAPI (blue). Representative images (C) and quantification (D) of 70 kDa FITC-dextran (white arrows) uptake in AsPC-1 cells pre-treated with EIPA (75 µM, 30 min) prior to EGF stimulation. Control cells received DMSO. Results represent mean ± SD of three independent experiments performed in triplicate. E. Quantification of 70 kDa FITC-dextran uptake in the indicated PDAC cells with or without EGF treatment (100 ng/mL, 5 min). 180 fields (∼25,000 cells) were analyzed per condition. Results represent mean ± SD of three independent experiments performed in triplicate. *p<0.05, **p<0.01, ***p<0.005, ****p<0.001.

### SG33 utilizes macropinocytosis for entry and replication in permissive PDAC cells

We investigated whether MPC contributes to SG33 entry in pancreatic cancer cells previously identified as permissive to SG33 infection (24). PDAC087T cells were infected with SG33 at a MOI of 16 and analyzed by immunofluorescence against viral proteins 6 hours post-infection. The MOI of 16 was selected to trigger a detectable infection using antibodies against viral proteins while avoiding premature cell lysis. Mock-infected cells served as controls. SG33 infection produced fluorescence predominantly localized in perinuclear complexes, consistent with viral replisomes (Fig. 3A, middle panel). Treatment with the MPC inhibitor EIPA reduced SG33 infection by 2.9-fold ± 1.2 (p<0.001, Fig. 3A, right panel; 3B), indicating that MPC mediates viral entry in PDAC087T cells.

**Figure 3.**
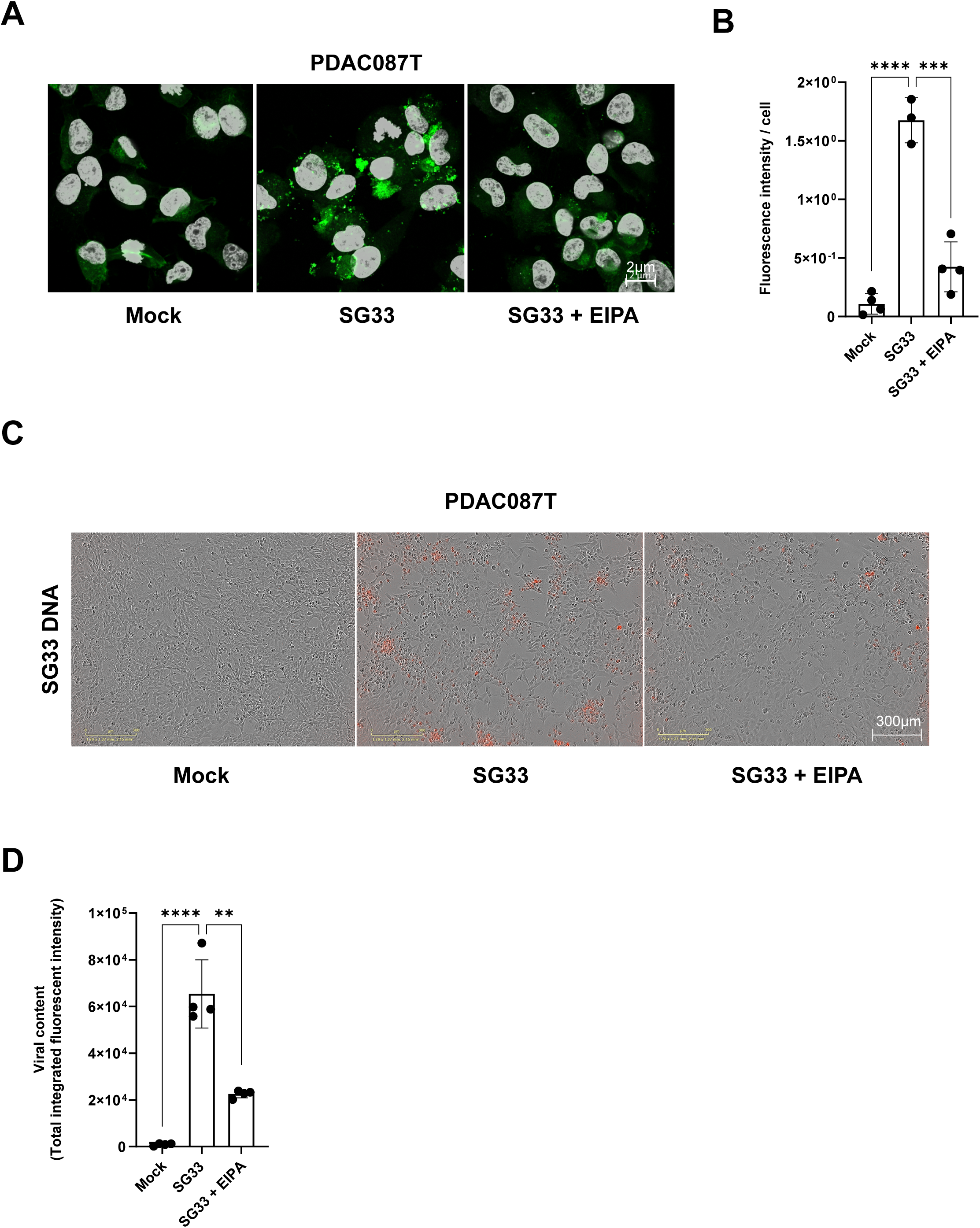
Macropinocytosis contributes to SG33 entry and replication. Representative images (A) and quantification (B) of SG33 entry in PDAC087T cells pre-treated with EIPA (50 µM, 30 min) followed by infection with SG33 (MOI 16). Control cells received DMSO. Approximately 5 fields (∼10 cells) were analyzed per condition. Results represent mean ± SD of four independent experiments. DAPI (grey) stains nuclei. Scale bars: 20 µm. Representative images (C) and quantification (D) of SG33-ANCHOR Santaka replication (MOI 5) in PDAC087T cells pre-treated with EIPA (25 µM, 30 min) and infected for 48 hours in the continuous presence of EIPA. Viral replication was quantified by integrated fluorescence intensity 72 h post-infection using IncuCyte Zoom. Results represent mean ± SD of four independent experiments performed in triplicate. *p<0.05, **p<0.01, ***p<0.005. Scale bars are indicated.

We previously demonstrated that the ANCHOR system allows real-time visualization of viral DNA in living cells by tagging the viral genome with a fluorescent reporter, enabling non-invasive monitoring of viral replication and intracellular localization (24). To assess the effect of MPC on viral replication, PDAC087T cells were infected with SG33 ANCHOR Santaka at a MOI of 5 and monitored for fluorescence non-invasively for up to 72 hours post-infection. Here again, the MOI of 5 was selected to trigger a detectable infection using the ANCHOR system without excessive cell lysis. Fluorescence for viral DNA increased in untreated, infected cells, reflecting active viral replication (Fig. 3C, middle panel), as compared to mock-infected control cells (Fig. 3C, left panel). EIPA treatment reduced viral replication by 64% ± 9 (p<0.01, Fig. 3C, right panel; 3D). These results demonstrate that MPC is required for efficient SG33 entry and replication in permissive PDAC cells.

### SG33 employs viral apoptotic mimicry to induce macropinocytosis and facilitate infection in PDAC cells

Previous studies have shown that viruses, including vaccinia virus, can trigger MPC to facilitate entry (20, 29). We investigated whether SG33 infection induces MPC in cells with inducible MPC. Serum-starved AsPC-1 cells were infected with SG33 at a MOI of 5, a condition that allows detection of infection without premature lysis, and analyzed for 70-kDa FITC-dextran uptake. As expected, EGF-treated AsPC-1 cells showed a 2.32-fold ± 0.9 increase in dextran uptake compared to mock-treated cells (p < 0.05, Fig. 4A right panel, B). Infection with SG33 increased dextran uptake 4.4-fold ± 1.65 (p < 0.05, Fig. 4A right panel, B). In PDAC087T primary cultures, EGF treatment increased dextran uptake 2.6-fold ± 1.25 relative to control cells (p < 0.05, Fig. 4C, D), whereas SG33 infection increased dextran uptake 9.5-fold ± 1.5 (p < 0.05, Fig. 4C, D). Treatment with EIPA reduced SG33-induced dextran uptake by 64% ± 15 (p < 0.05, Fig. 4C, D), confirming that SG33 induces MPC. To exclude non-specific effects, PDAC051T cells (permissive to SG33 but displaying low basal and inducible MPC) were infected with SG33 at a MOI of 5 and incubated with FITC-dextran. Under these conditions, no detectable increase in dextran uptake was observed (Fig. 4E). Collectively, these data indicate that SG33 infection induces MPC in PDAC primary cells.

**Figure 4.**
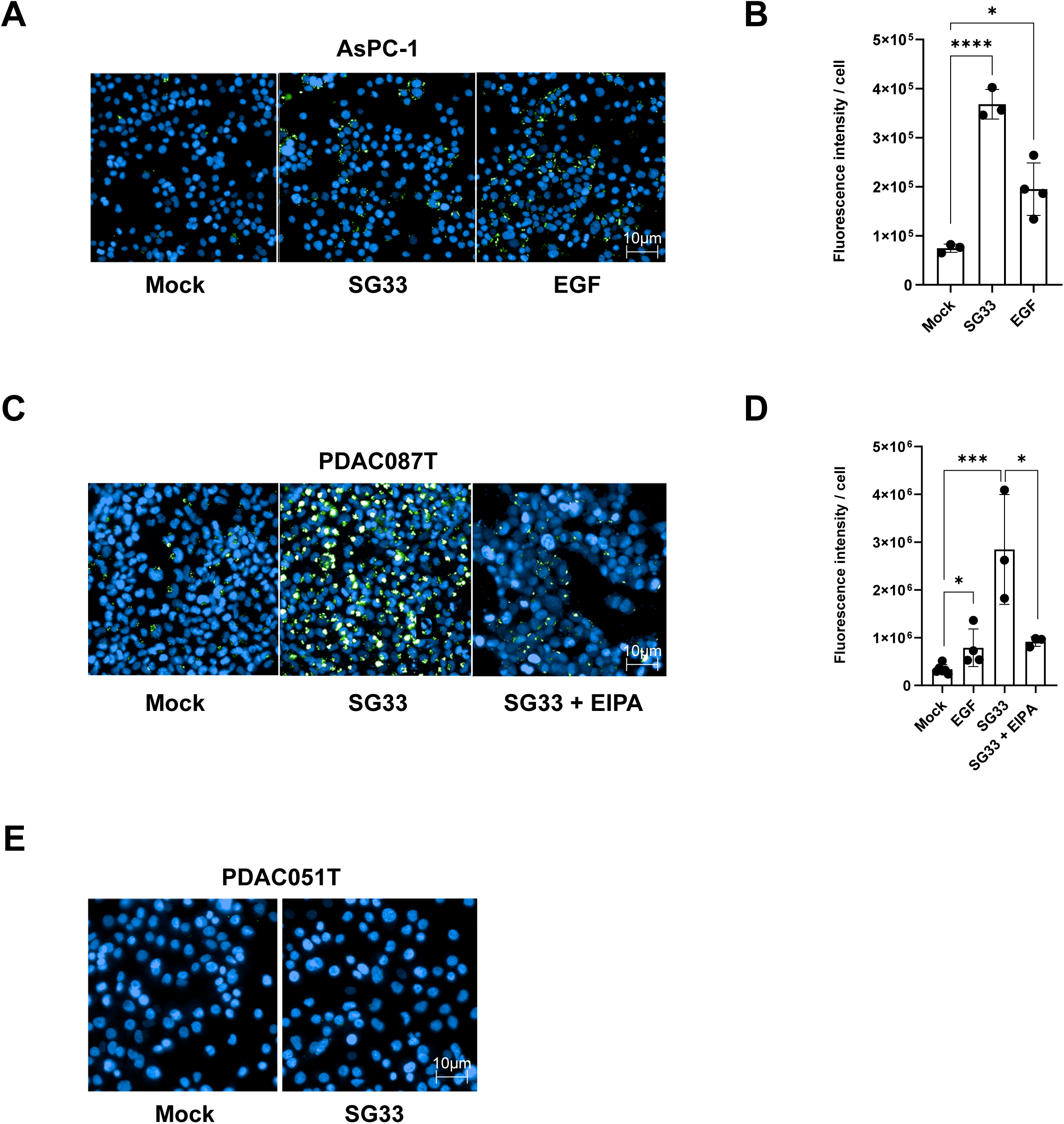
SG33 stimulates macropinocytosis in a subset of PDAC cells. Representative images (A) and quantification (B) of 70 kDa FITC-dextran uptake in AsPC-1 cells following SG33 infection (MOI 5). EGF (100 ng/mL, 5 min) serves as a positive control. Control cells were cultured with 2 mM glutamine. Results represent mean ± SD of at least three independent experiments performed in triplicate. Nuclei stained with DAPI (blue). Representative images (C) and quantification (D) of 70 kDa FITC-dextran uptake in PDAC087T cells pre-treated or not with EIPA (75 µM, 30 min) and infected or not with SG33 (MOI 5). EGF treatment serves as positive control. 180 fields (∼25,000 cells) were analyzed per condition. Results represent mean ± SD of three independent experiments performed in triplicate. E. Representative images of 70 kDa FITC-dextran uptake in PDAC051T cells with or without SG33 infection (MOI 5). Representative of three independent experiments. *p<0.05, ***p<0.005. Scale bars are indicated.

We next investigated the molecular mechanism of SG33-induced MPC. Vaccinia virus (MV form) induces MPC via apoptotic mimicry, exposing phosphatidylserine on the viral membrane (20). To determine whether SG33 employs a similar mechanism, SG33 ANCHOR Santaka particles were sonicated for single-virion analysis, as it is known that the ANCHOR system allows visualization of large viruses even outside cells (30). Viral suspensions were incubated with fluorescent Annexin-V to detect phosphatidylserine. The vast majority (91%) of SG33 particles were Annexin-V positive (Fig. 5A left panel). Treatment with Nonidet-P40 to remove phospholipids abolished Annexin-V binding (Fig. 5A right panel), confirming phosphatidylserine presence on the viral membrane. To assess functional relevance, SG33 particles were pre-incubated with Annexin-V before infection. Serum-starved PDAC087T cells were infected with untreated SG33 or Annexin-V-saturated SG33 at a MOI of 5. Uptake of 70-kDa FITC-dextran was reduced 71% ± 33 in cells infected with Annexin-V-treated SG33 (p<0.05, Fig. 5B, C). As expected, viral protein expression 6 hours post-infection showed perinuclear localization in untreated, SG33-infected cells (Fig. 5D, middle panel). Pre-incubation with Annexin-V reduced fluorescent signal by 79% ± 23 (p<0.01, Fig. 5D, right panel, 5E). These results indicate that phosphatidylserine on the SG33 envelope contributes to MPC induction and facilitates viral entry and infection in cells capable of inducible MPC.

**Figure 5.**
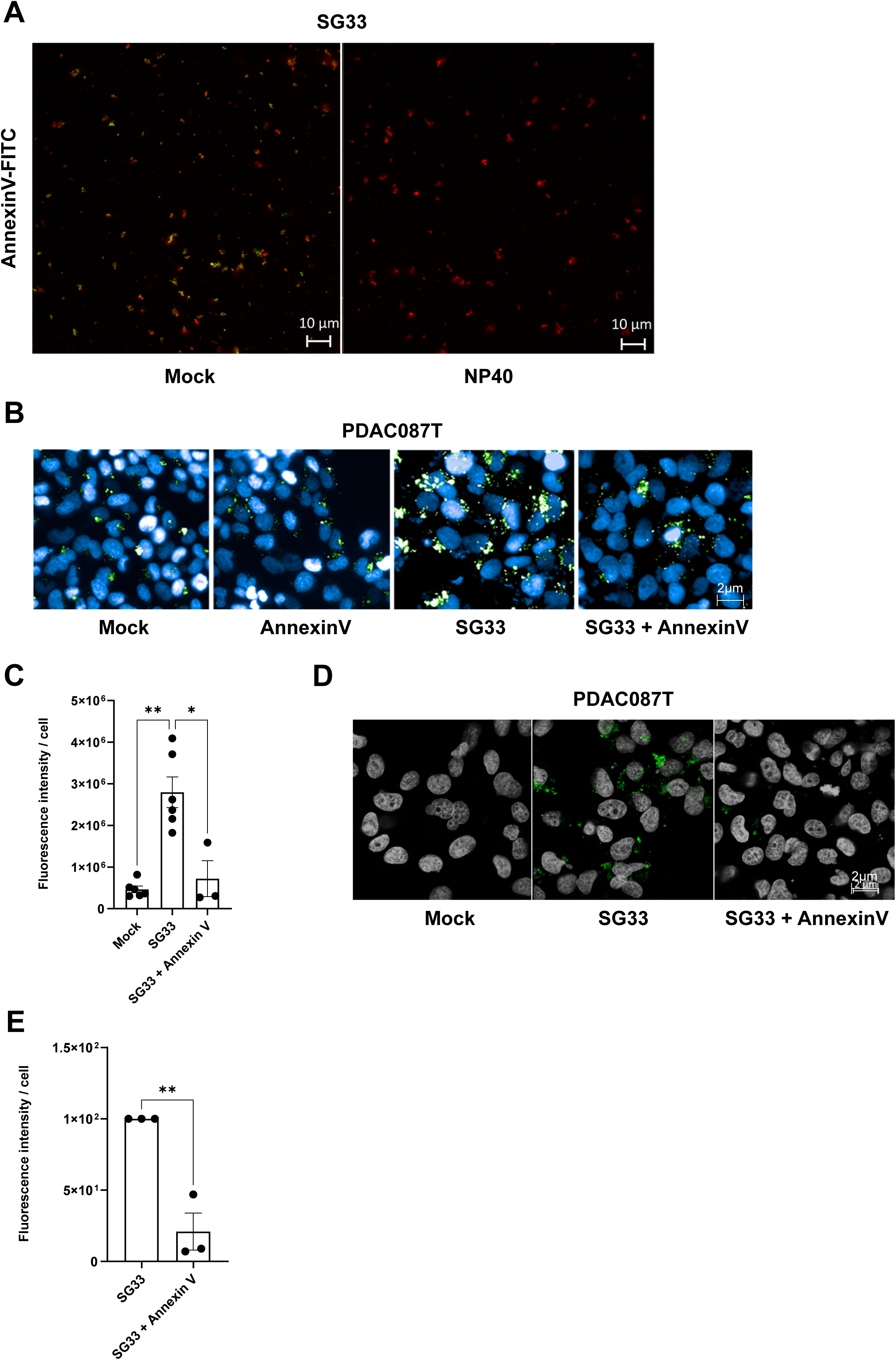
Role of viral phosphatidylserine in SG33-induced macropinocytosis. (A) SG33 ANCHOR Santaka particles (red) were sonicated and treated or not with 0.5% NP-40 for 30 min, followed by 15 min incubation with AnnexinV-FITC. Pictures presented are representative of three experiments. Representative images (B) and quantification (C) of 70 kDa FITC-dextran uptake in PDAC087T cells pre-treated or not with AnnexinV (15 min) and infected or not with SG33 (MOI 5). Representative images (D) and quantification (E) of SG33 infection (MOI 5) in PDAC087T cells pre-treated or not with AnnexinV. 5 fields (∼10 cells) were analyzed per condition. Results represent mean ± SD of three independent experiments performed in triplicate. DAPI (blue) stains nuclei. *p<0.05, **p<0.01. Scale bars are indicated.

### SG33 utilizes alternative endocytic pathways to infect PDAC cells

Our results identified MPC as an important mechanism for SG33 infection in PDAC cells. However, some SG33-permissive cells exhibit low basal MPC or lack inducible MPC (24). We therefore investigated alternative endocytic pathways that SG33 might exploit. Clathrin-mediated endocytosis is a well-characterized route used by many viruses for cellular entry(6). To test its role in SG33 infection, PDAC087T cells were treated with hydroxy-dynasore, a dynamin inhibitor, in a dose-response manner. Transferrin uptake was quantified as a marker of clathrin-mediated endocytosis. Treatment with 400 μM hydroxy-dynasore inhibited transferrin uptake by 86% ± 8 (p<0.05, Fig. 6A, B). These conditions were used for subsequent experiments. PDAC087T cells were infected with SG33 at a MOI of 16 in the presence or absence of hydroxy-dynasore. Viral entry was analyzed by immunofluorescence 6 hours post-infection, with mock-infected cells as controls. In untreated cells, fluorescence indicative of viral replisomes localized near the nucleus (Fig. 6C, top panel and focus, white arrow). Quantification showed that 43% ± 2 of cells exhibited perinuclear replisomes, compared to 8% ± 2 in hydroxy-dynasore-treated cells, corresponding to an 80% ± 5 reduction (p<0.001, Fig. 6D). In the presence of hydroxy-dynasore, viral signal remained at the cell periphery (Fig. 6C, bottom panel and focus, grey arrow). We further pursued these investigations using PDAC051T cultures, which are permissive to the virus (24) but in which we failed to detect macropinocytic activity (Fig. 1D and 2E). These cells were treated with hydroxy-dynasore and subsequently infected with SG33 at an MOI of 5. The results shown in Figure 6E–F demonstrate that hydroxy-dynasore treatment significantly inhibits viral entry into these cells. Indeed, endocytosis inhibition reduced by 82% the number of cells displaying viral replisomes (Fig. 6F, p<0.001). These data indicate that SG33 also exploits clathrin-mediated endocytosis to infect PDAC cells lacking robust MPC activity.

**Figure 6.**
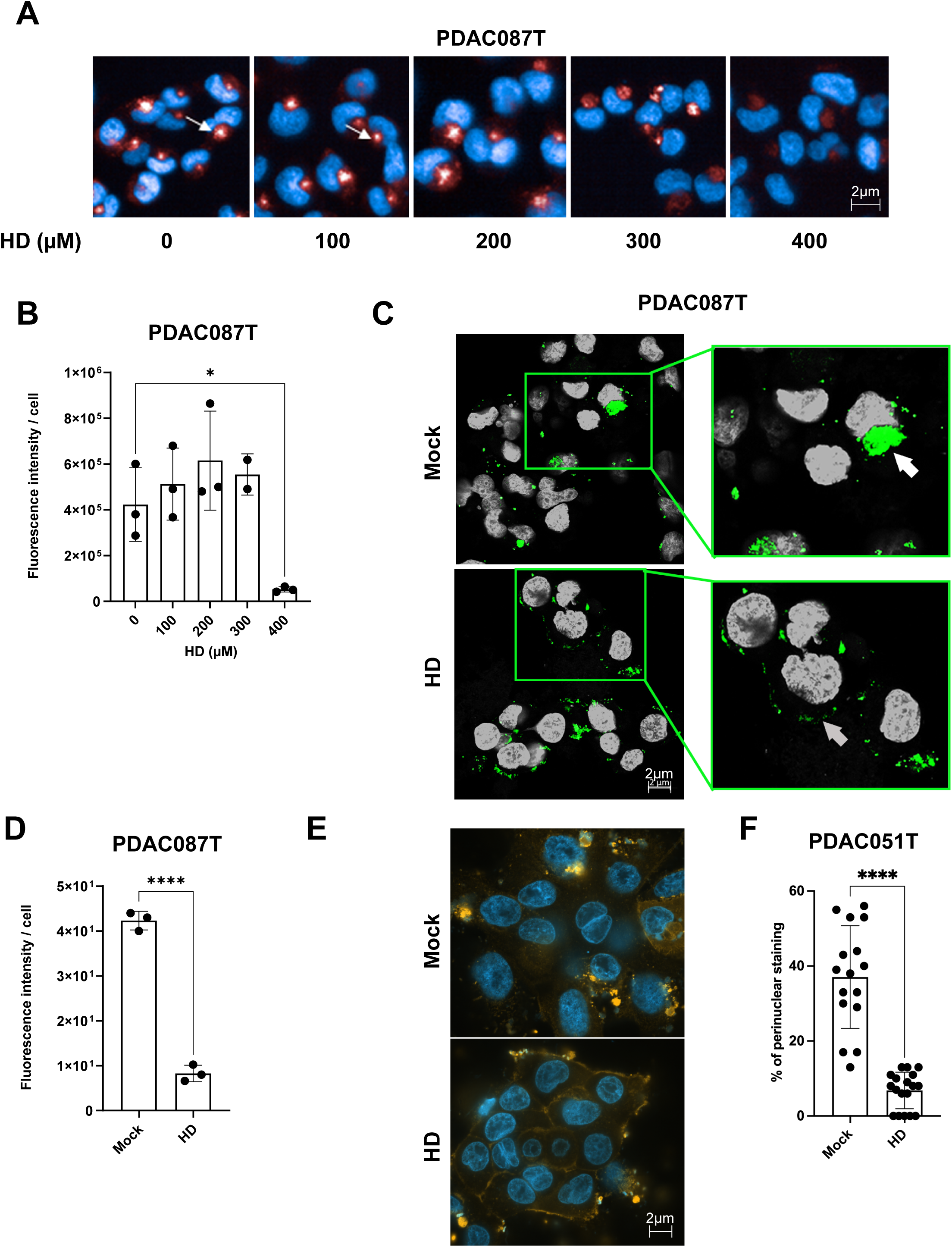
SG33 uses clathrin-mediated endocytosis in PDAC cells. Representative images (A) and quantification (B) of transferrin uptake in PDAC087T cells treated with hydroxy-dynasore (HD) at indicated doses for 2h. Control cells received DMSO. Approximately 180 fields (∼25,000 cells) were analyzed per condition. Results represent mean ± SD of three independent experiments performed in triplicate. Nuclei stained with DAPI (blue). Representative images (C) and quantification (D) of SG33 infection (MOI 16) in PDAC087T cells pre-treated with HD (400 µM, 2h). White arrows indicate SG33 foci near nuclei (grey); grey arrows indicate SG33 foci at the plasma membrane. 5 fields (∼10 cells) were analyzed per condition. Representative of three experiments. *p<0.05, ****p<0.001. Scale bars are indicated. PDAC051T cells were infected with SG33 at MOI=5 in the presence or not of HD (200µM, 2h). Representative images (E) and quantification (F) of SG33 infection. 6 fields (175-236 cells) were analyzed per condition. Representative of three experiments. ****p<0.001. Scale bars are indicated.

## DISCUSSION

Oncolytic virotherapy has advanced considerably, yet no OV has been approved for PDAC, one of the most lethal malignancy. Viral tropism, defined as the ability of viruses to infect specific cell types, is a critical determinant of therapeutic efficacy. A deeper understanding of entry mechanisms may help identify PDAC subtypes more permissive to infection and guide the development of targeted therapies.

Viruses typically enter cells either through direct fusion at the plasma membrane or via endocytosis (6). Endocytic uptake provides several advantages, including bypassing cytoskeletal barriers, releasing genomes at defined intracellular sites, protecting viral proteins from immune recognition, and shielding fusion proteins from neutralizing antibodies(31). Among these pathways, MPC is characterized by the actin-driven formation of large, uncoated vesicles and non-selective uptake of extracellular fluid(9). MPC is frequently dysregulated in cancer, where it contributes to nutrient scavenging and supports the metabolic demands of tumor cells (9).

Building on evidence that vaccinia virus triggers MPC for entry (20) and that MYXV may exploit an MPC-like route(22), we investigated whether the MYXV attenuated SG33 strain engages MPC in PDAC cells. By characterizing basal and inducible MPC across cell lines and patient-derived cultures, we confirmed previously reported differences (16, 27) (low basal MPC in BxPC-3, high in MIA PaCa-2, inducible MPC in AsPC-1 cells). We extended this characterization to primary PDAC cultures, despite the fact that primary cells often do not tolerate glutamine starvation, likely due to the critical role of glutamine in supporting PDAC cell proliferation and redox balance (32). This also may explain the decrease observed in dextran uptake in PDAC087T starved for this amino acid. However, most patient-derived cultures exhibited low basal MPC with limited inducibility, underscoring a greater heterogeneity than previously appreciated and suggesting that MPC prevalence in PDAC may have been overestimated.

We next examined whether MPC contributes to SG33 infection. Pharmacological inhibition impaired both entry and replication, supporting a functional role for MPC. This is consistent with reports that MPC facilitates entry of diverse viruses, including adenoviruses 3 and 35 (33), echovirus 1, Ebola virus (34), influenza A virus, (35) measles virus (36), Nipah virus (37), respiratory syncytial virus (38), and HIV (39). MPC provides distinct advantages for viral entry, enabling internalization of particles too large for other endocytic routes and facilitating infection of a broader range of host cells and tissues. This is largely due to its non-specific nature, which allows viruses to exploit the pathway without requiring engagement of specific entry receptors (8). Our findings are in contrast with a previous study that proposed MYXV and vaccinia virus utilize distinct entry pathways in human cancer cells (21). In that work, MPC involvement was excluded on the basis of experiments using genistein; however, this conclusion is questionable given the limited specificity of genistein as an inhibitor. Moreover, results shown involving PAK1 status, a central regulator of MPC, leave some uncertainties regarding the interpretation of MPC involvement.

At the mechanistic level, we show that SG33 induces MPC in permissive PDAC cells and that viral membrane-associated phosphatidylserine contributes to this induction, consistent with apoptotic mimicry described for vaccinia virus (20). These observations extend prior evidence implicating phosphatidylserine in poxvirus infectivity (40, 41) and in Zika virus transmission, where extracellular vesicle-associated phosphatidylserine competed with viral entry (29). Our results suggest that SG33 harnesses a similar mechanism to trigger MPC. Importantly, virus-induced MPC may also be exploited therapeutically. For instance, IGF1R inhibitors stimulate MPC and enhance delivery of nanoparticulate albumin-bound paclitaxel (42). It will be of interest to determine whether SG33-induced MPC similarly potentiates the uptake and activity of clinically relevant MPC-dependent drugs.

In PDAC cells with low or absent MPC, SG33 infection remained efficient, and our data indicate that clathrin-mediated endocytosis can serve as an alternative entry route. Such redundancy parallels strategies used by other viruses, such as Puumala virus (43), which flexibly exploits multiple endocytic pathways depending on host context. This adaptability likely ensures infection across heterogeneous tumor populations and under variable microenvironmental conditions. Overall, our study identifies MPC as a virus-induced entry route for SG33 in PDAC cells and demonstrates that clathrin-mediated endocytosis provides an alternative pathway when MPC is limited. This dual entry strategy highlights the flexibility of SG33 and reinforces its potential as a versatile oncolytic agent for PDAC.

## MATERIALS AND METHODS

### Reagents

5-(N-Ethyl-N-isopropyl) amiloride hydrochloride (EIPA; HY-101840A) was purchased from Clinisciences. FITC-dextran (10 kDa, 11520226; 70 kDa, 11500226) and transferrin (11550766) were from Thermo Fisher Scientific. Epidermal growth factor (EGF; 17883153) was from Peprotech. Annexin-V-FITC (556547) and Annexin-V (556416) were from BD Biosciences. Nonidet-P40 (74385-1L) was from Sigma-Aldrich. Hydroxy-dynasore (5364/10) was from Bio-Techne.

### Viral cloning, production and titration

The recombinant virus was generated by homologous recombination using a donor plasmid derived from pSK-Vmyxlac, designed to target the intergenic region between M009L and M010 of the SG33 genome, as previously described(24). The recombination cassette contained the ANCHOR3 system (NeoVirTech) fused to a guanine phosphoribosyltransferase of E. coli origin (*ecoGPT*) selectable marker to allow positive selection of recombinant viruses. The ANCHOR3-Santaka cassette was placed under the control of the viral pMH5 promoter to ensure efficient expression during infection. Santaka is a synthetic monomeric red fluorescent protein (λ_ex 551 nm; λ_em 571 nm), engineered for high photostability and brightness, making it suitable for real-time visualization of viral DNA in living cells (25). For viral production, 3.5 × 10⁶ RK13 cells were grown to confluence in T175 flasks (Thermo Fisher Scientific; 178883) in complete DMEM. Cells were washed once with PBS and infected with SG33 at a multiplicity of infection (MOI) of 0.5 at 37°C. Following infection with either SG33 or SG33-ANCHOR-Santaka, cells were incubated for 72 h at 37°C until complete cytopathic effect was observed. Cells were then frozen at –20°C overnight. The crude supernatant containing viral particles was collected and stored at –20°C. Viral titers were determined by plaque assay. Briefly, 9 × 10³ RK13 cells were seeded in 96-well plates (Corning) in complete DMEM and incubated for 24 h at 37°C. Virus was serially diluted from 10⁻¹ to 10⁻¹¹ and added to the cells for 1 h at 37°C. The inoculum was then replaced with complete DMEM. Viral plaques were counted 48 h post-infection from dilutions 10⁻³ to 10⁻⁶.

### Cell culture

AsPC-1 (ATCC-CRL-1682) and BxPC-2 (ATCC-CRL-1687) cells were cultured in RPMI (GlutaMAX; Life Technologies) supplemented with 1% penicillin/streptomycin (P/S; Life Technologies). MIA-PaCa-2 (ATCC-CRL-1420), RK13(24), PDAC012T, PDAC015T, PDAC084T, PDAC087T, and PDAC091T cells were cultured in DMEM (4.5 g/L glucose; Life Technologies) with 1% P/S, as previously described(26). PDAC051T cells were grown in serum-free ductal medium (SFDM) as previously described. All cell lines except PDAC051T were supplemented with 10% fetal bovine serum (FBS; Thermo Fisher Scientific). Hereafter, these media are referred to as complete media.

Cells were maintained at 37°C with 5% CO₂ and routinely tested for mycoplasma contamination (Invivogen, Mycoplasma detection kit). For serum starvation experiments, cells were cultured in serum-free medium for 18 h (0% FBS). PDAC051T cells were starved for 3 h in DMEM/F12 (Life Technologies) supplemented with 10 mM nicotinamide and 28 mM D-glucose. For glutamine starvation, cells were cultured in glutamine-free RPMI or DMEM (Life Technologies) without serum. For PDAC051T cells, glutamine-free DMEM/F12 (Life Technologies) supplemented with 10 mM nicotinamide and 28 mM D-glucose was used. All starvation media were phenol-red free (to minimize autofluorescence) and supplemented with 1% P/S.

### Monitoring and quantification of dextran uptake

Cells were seeded in black, optically clear, flat-bottom 96-well plates (6055302, Perkin Elmer) in complete media and incubated for 24 h. Seeding densities were adjusted for each cell line to reach ∼80% confluence at the time of the experiment: MiaPaCa-2, AsPC-1, BxPC-3, and PDAC012T (1.5 × 10⁴ cells/well); PDAC087T, PDAC084T, PDAC015T, PDAC072T, and PDAC051T (1.75 × 10⁴ cells/well); PDAC091T (2 × 10⁴ cells/well). Cells were subjected to serum or glutamine starvation as described above. FITC-dextran (10 or 70 kDa) was added to starvation media at 1 mg/ml for 30 min at 37°C. For inhibition or induction of macropinocytosis (MPC), cells were pre-treated with 75 μM EIPA for 30 min or 100 ng/ml EGF for 5 min, respectively, prior to FITC-dextran addition. For infection assays, serum-starved cells were infected with SG33 (3.1 × 10⁶ pfu/ml; MOI 5) for 30 min at 4°C to allow viral attachment. Cells were then incubated with 70 kDa FITC-dextran (1 mg/ml, 30 min, 37°C), washed three times with ice-cold PBS, and fixed in 4% paraformaldehyde (15710-5, Electron Microscopy Sciences). Nuclei were counterstained with DAPI (1050A, Euromedex). Fluorescence images were acquired using the Operetta CLS high-content analysis system (Revvity), equipped with an automated 40×/0.6 NA air objective. Excitation was performed with high-power LEDs at 355–385 nm (DAPI), and 460–490 nm (FITC, with emission collected through appropriate filters (e.g., 430–500 nm for DAPI, and 500–550 nm FITC). Fluorescence quantification was performed with Harmony software (v4.9). Digital Phase Contrast was used to delineate cytoplasmic boundaries. Nuclei, cytoplasm, and FITC-dextran spots were identified using the “Find Nuclei,” “Find Cytoplasm,” and “Find Spots” modules. Spots within the size range of macropinosomes (0.2–5 μm) were selected. The macropinocytosis index was calculated as the total fluorescence intensity per cell.

### Western blotting

AsPC-1 cells were seeded in 100-mm culture dishes (Corning) at 2.5 × 10⁶ cells/dish in 10 mL of complete medium and incubated for 24 h at 37°C. After two PBS washes, cells were subjected to serum deprivation for 24 h. EGF (25 ng/mL) was added directly to the medium for 5, 15, or 30 min at 37°C. Cells were washed twice with ice-cold PBS, and proteins were extracted on ice using RIPA buffer (50 mM Tris-HCl pH 7.4, 150 mM NaCl, 1% Nonidet-P40, 1% sodium deoxycholate, 1% SDS) supplemented with protease inhibitors (P8340, 1X, Sigma) and phosphatase inhibitors (1861277, 1X, Thermo Fisher Scientific). Cell lysates were sonicated (10 s, 40% amplitude; Vibra-Cell) and centrifuged (16,049 × g, 15 min, 4°C).

Protein concentration was determined with the Bio-Rad Protein Assay Dye Reagent (5000006, Bio-Rad). Equal amounts of protein (40 μg) were denatured in Laemmli buffer (95°C, 5 min), resolved by SDS-PAGE (7.5–10%), and transferred to nitrocellulose membranes (Bio-Rad) using the TransBlot Turbo system. Membranes were blocked with 10% BSA (04100812E, Euromedex) in TBS-0.1% Tween-20, then incubated overnight at 4°C with primary antibodies: p-EGFR (Y1068, 3777, 1:1000, CST), EGFR (4267, 1:1000, CST), p-PAK1 (Ser144)/PAK2 (Ser141, 2606, 1:500, CST), α-PAK (sc-166887, 1:500, Santa Cruz), p-ERK1/2 (Thr202/Tyr204, 4370, 1:1000, CST), ERK1/2 (4695, 1:2000, CST), β-ACTIN (3700, 1:1000, CST). After three washes, membranes were incubated with HRP-conjugated secondary antibodies (1:10,000, 1 h, RT). Proteins were detected using Clarity Western ECL substrate (Bio-Rad) and imaged on a Chemi-Doc CRS+ system (Bio-Rad). For phosphorylation analysis, membranes were first probed with phospho-specific antibodies, stripped with Restore PLUS buffer (46430, Thermo Fisher Scientific), and re-probed with antibodies against total proteins.

### Immunofluorescence for viral proteins

PDAC087T cells were seeded in 8-well Lab-Tek slides (154941, Thermo Fisher Scientific) at 2 × 10⁴ cells/well in 200 μL of complete medium and incubated for 24 h at 37°C. Cells were treated with EIPA (50 μM, 30 min, 37°C) prior to infection with SG33 (MOI 16, 1 h, 37°C). After washing with PBS, cells were incubated in complete DMEM for 5 h, then fixed with 4% paraformaldehyde (15 min, RT) and permeabilized with 0.2% Triton X-100 (15 min). After blocking with 10% BSA (2 h, RT), cells were incubated overnight at 4°C with a rabbit anti-SG33 primary antibody (1:500; kind gift from Prof. S. Bertagnoli, ENVT) in 3% BSA. Cells were washed and incubated with Alexa Fluor-488 anti-rabbit IgG (A11008, 1:1000, Thermo Fisher Scientific) for 2h at RT. Nuclei were counterstained with DAPI (1:10,000, 5 min, RT). Slides were mounted in Fluorescent Mounting Medium (S3023, DAKO). Imaging was performed on a Zeiss 880 Fast Airyscan confocal microscope with a 63× oil-immersion objective (NA 1.4). Excitation was performed using diode lasers at 405 nm, and 488 nm, and emission was collected using standard filter sets. The Airyscan detector allowed super-resolution imaging with improved signal-to-noise ratio. Images were analyzed with Fiji (v2.1.0-1) using macro-assisted analysis.

### Real time monitoring of viral replication in cell populations

PDAC087T cells were seeded in 96-well plates (Corning) at 1 × 10⁴ cells/well in 100 μL complete medium. After 24 h, cell counts were used to adjust MOI. Cells were washed with PBS and infected with SG33-ANCHOR-Santaka (5 × 10⁶ pfu/mL; MOI 5) for 1 h at 37°C in phenol-red–free medium. Medium was then replaced with fresh phenol-red–free complete medium. Real-time imaging was performed using the IncuCyte Zoom system (Sartorius) at 37°C, 5% CO₂. Images were acquired every 3 h at 10× magnification in bright-field and red fluorescence channels (Ex 565–605 nm, Em 625–705 nm). Data were analyzed with Zoom2018A software.

### Imaging and analysis of viral particles

To visualize individual particles, SG33-ANCHOR-Santaka was sonicated (30 s, 25% amplitude, 25 W) and centrifuged (406 × g, 10 min). A total of 1 × 10⁴ particles (1.9 × 10⁵ pfu/mL) were seeded on 18-well IBIDI slides (81817, Clinisciences). Fluorescent particles were imaged using a Zeiss 880 Fast Airyscan confocal microscope with a 63× oil-immersion objective (NA 1.4). Spot size was quantified using Fiji macros. Spot areas (µm²) were converted to radii (nm) using the equation: R = √(area/π).

### Visualization of phosphatidylserine at the viral membrane

For Annexin-V staining, 1 × 10⁴ SG33-ANCHOR-Santaka particles (1.9 × 10⁵ pfu/mL), previously sonicated (30 s, 25% amplitude, 25 W), were incubated with 54 ng Annexin-V-FITC (556547, BD Biosciences) in Annexin binding buffer (0.1 M HEPES-NaOH pH 7.4, 1.4 M NaCl, 25 mM CaCl₂) and phenol-red–free DMEM (final volume 60 µL). Samples were incubated for 15 min at RT, protected from light, and seeded on 18-well IBIDI slides. Colocalization between red (Santaka) and green (Annexin-V-FITC) signals was quantified with Fiji macros. For delipidation assays, sonicated SG33-ANCHOR-Santaka particles were treated with 0.5% Nonidet-P40 (30 min, RT), then stained with Annexin-V-FITC as above. Imaging was performed with the Zeiss 880 Fast Airyscan microscope.

### Statistical analysis

All statistical analyses were performed using GraphPad Prism v10.1.2. Unless otherwise stated, experiments were conducted at least in triplicate. Results are presented as mean ± SD. Statistical significance was assessed using unpaired two-tailed Student’s t-tests or one-way ANOVA. p values < 0.05 were considered significant (p<0.05; *p<0.01; **p<0.001; ***p<0.0001).

## Acknowledgements

We thank Damien Varry (Cancer Research Center of Toulouse; CRCT) for help with the Fiji macro command and Dr. J. Enninga (Pasteur Institute, Paris, France) for advising, N. Hanoun-Ziane, E. Martin and C. Zanibellato (CRCT) for administrative support. We also thank the technological pole of CRCT.

## Authors contributions

NK: conceptualization, data acquisition and analysis, writing–original draft. ND: conceptualization, data acquisition and analysis. LL: conceptualization, data acquisition and analysis. ST, FG and SB: conceptualization and production of viral strains. ND: conceptualization, writing–review and editing. LB: conceptualization, supervision, writing–review and editing. PC: conceptualization, data curation, data analysis, supervision, funding acquisition, validation, methodology, writing–review and editing.

