## supplementary figure 1 for "SG33, a vaccine strain of myxoma virus with oncolytic potential, exploits macropinocytosis and clathrin-mediated endocytosis for entry into pancreatic cancer cells"

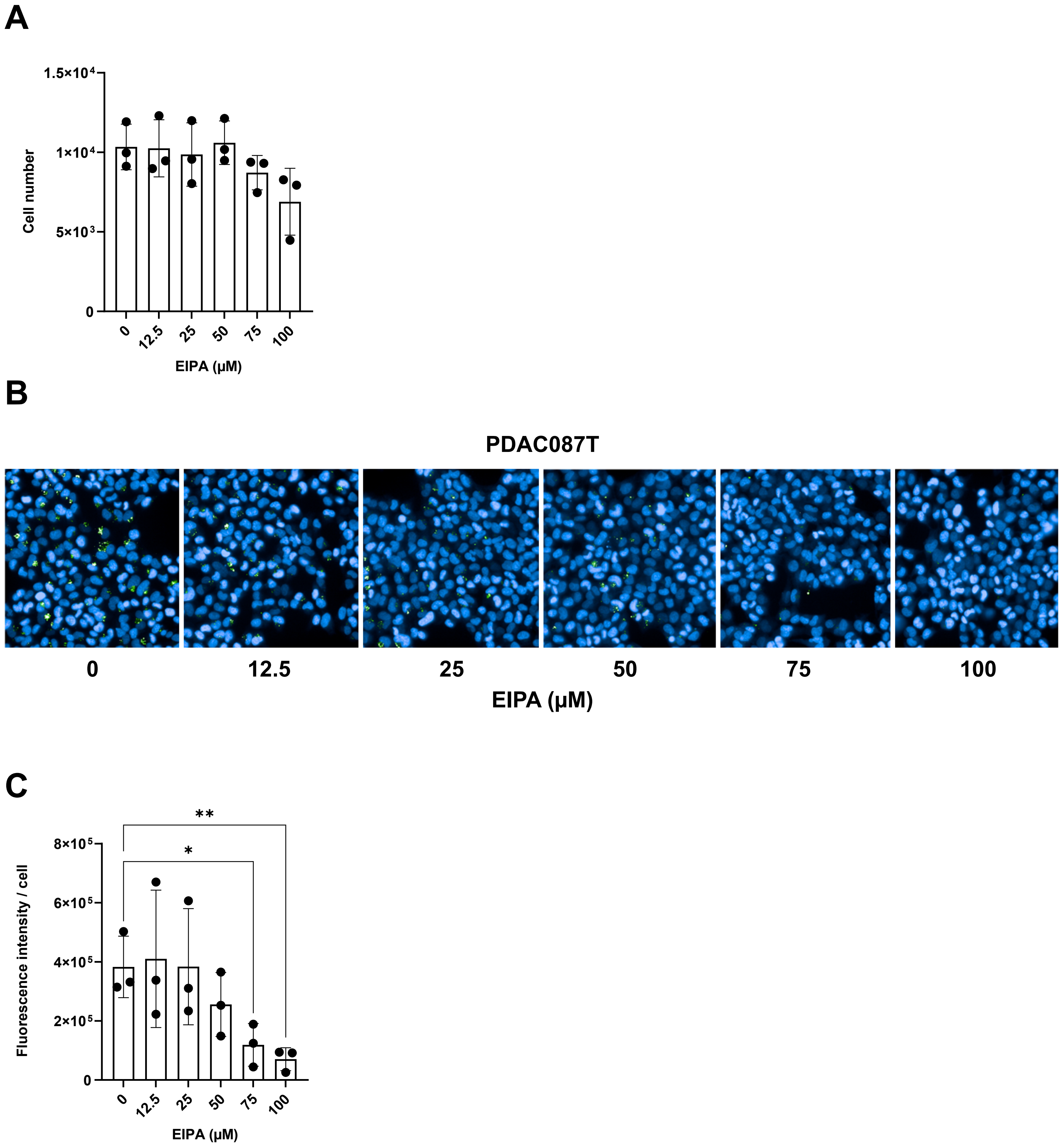


Supplemental Figure 1. Dose-response effect of EIPA on PDAC087T cells. (A) Nuclei count following EIPA treatment for 30 min. Approximately 180 fields (~25,000 cells) analyzed per condition. Results represent mean ± SD of three independent experiments. (B, C) Representative images and quantification of 70 kDa FITC-dextran uptake in PDAC087T cells after EIPA treatment. Control cells received DMSO. DAPI (blue) stains nuclei. *p<0.05, **p<0.01. Scale bars indicated.
