## supplementary figure 2 for "SG33, a vaccine strain of myxoma virus with oncolytic potential, exploits macropinocytosis and clathrin-mediated endocytosis for entry into pancreatic cancer cells"

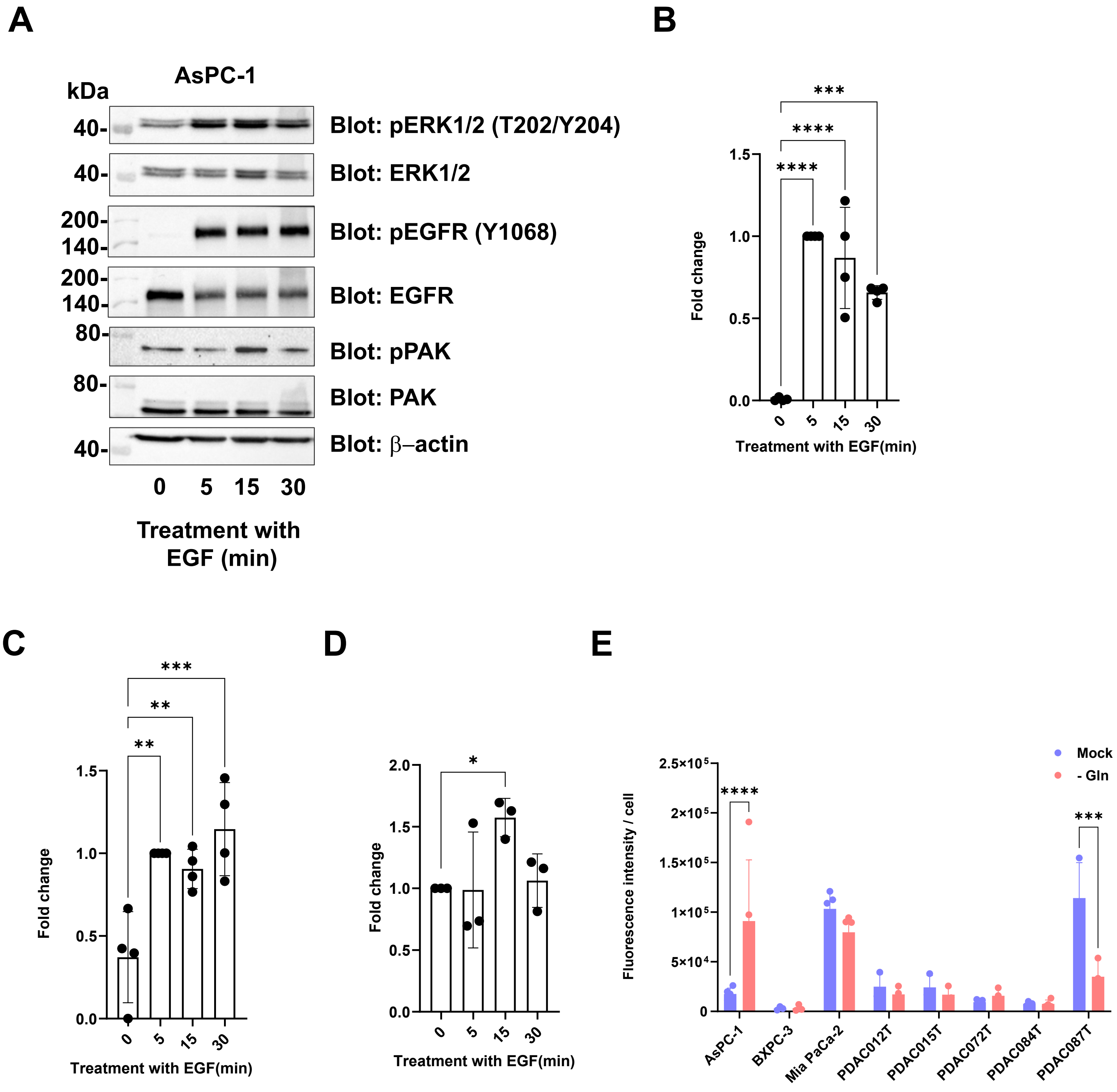


Supplemental Figure 2. EGFR, ERK1/2, and PAK activation in AsPC-1 cells, and characterization of inducible macropinocytosis in PDAC cells. (A) Representative immunoblots and (B–D) quantification of EGFR, ERK1/2, and PAK phosphorylation following EGF treatment (25 ng/mL) at indicated times. β-actin used as loading control. (D) Quantification of 70 kDa FITC-dextran uptake in the presence or absence of 2 mM glutamine. Results represent mean ± SD of at least three independent experiments performed in triplicate. *p<0.05, **p<0.01, ***p<0.005, ****p<0.001.
